# Halyos: A patient-facing visual EHR interface for longitudinal risk awareness

**DOI:** 10.1101/597583

**Authors:** Samson Mataraso, Vimig Socrates, Fritz Lekschas, Nils Gehlenborg

**Affiliations:** Harvard Medical School, Department of Biomedical Informatics; University of California, Berkeley, Department of Electrical Engineering and Computer Sciences; University of California, Berkeley, Department of Bioengineering; Case Western Reserve University, Department of Electrical Engineering and Computer Science; Harvard John A. Paulson School of Engineering and Applied Sciences

**Author notes:** = these authors contributed equally to this work (Samson Mataraso, Vimig Socrates). = corresponding author.

**Keywords:** Data Interpretation, Data Visualization, Electronic Health Records, Patient Portal, Risk Management

## Abstract

We have developed Halyos (http://halyos.gehlenborglab.org), a visual EHR web application that complements the functionality of existing patient portals. Halyos is designed to integrate with existing EHR systems to help patients interpret their health data. The Halyos application utilizes the SMART on FHIR (Substitutable Medical Applications and Reusable Technologies on Fast Healthcare Interoperability Resources) platform to create an interoperable interface that provides interactive visualizations of clinically validated risk scores and longitudinal data derived from a patient’s clinical measurements. These visualizations allow patients to investigate the relationships between clinical measurements and risk over time. By enabling patients to set hypothetical future values for these clinical measurements, patients can see how changes in their health will impact their risks. Using Halyos, patients are provided with the opportunity to actively improve their health based on increased understanding of longitudinal information available in EHRs and to begin a dialogue with their providers.

## INTRODUCTION

Patients are gaining greater access to data stored in Electronic Health Records (EHRs) through patient-facing personal health records (PHRs). However, many patients have difficulty understanding their own health information [1],[2]. Properly aggregating, representing, and explaining these data has great potential to empower patients in making informed decisions about their health, promote preventive care, and reduce burden on physicians by providing patients with an alternate source of personalized, interpretable health information. We introduce Halyos (Figure 1) as a web-based health exploration application that allows patients to explore their health data through clinically validated risk scores for disease, visualize their longitudinal data, and understand how changes in their health (e.g., increased blood pressure) impact their future disease risk scores.

**Figure 1.**
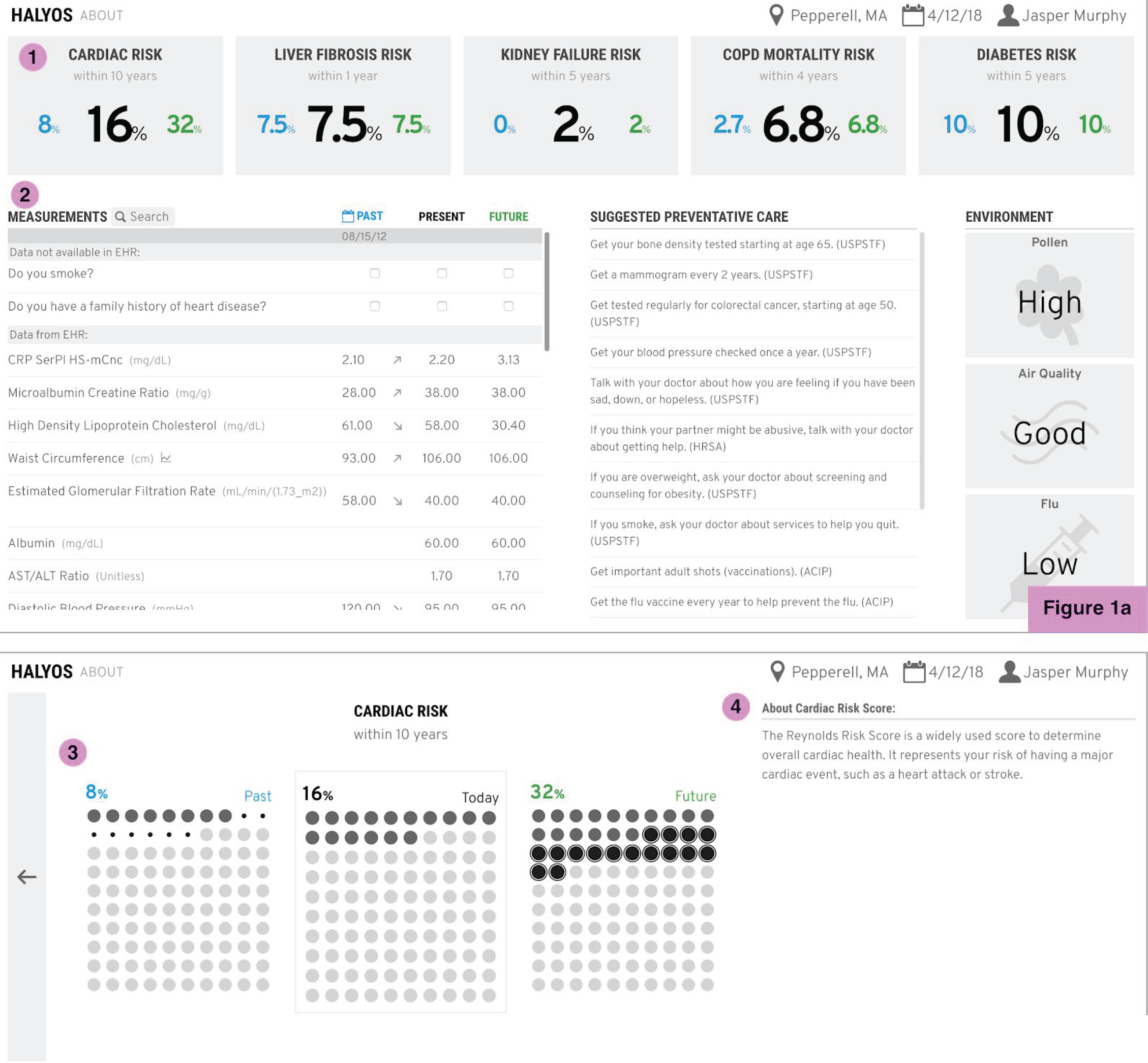
(1a, top): Home screen of Halyos. Risk scores are calculated from patient measurements and displayed as percentages (1). Individual measurement values are displayed under the Measurements section (2) with a detailed view shown in Figure 2. The dashboard also includes suggestive preventative care and environmental factors affects health risks. (1b, bottom): Expanded risk tile which displays pictorial unit charts representing a patient’s past, present, and future risk (3) with a written description of the risk score (4).

Although nearly 90% of hospitals provide the ability for patients to view, download, and transmit health records (<u>Office of the National Coordinator for Health Information Technology</u>), they cannot be used effectively if patients struggle to understand the information presented to them [1,2]. As they exist presently, patient portals do not improve health outcomes, in part due to comprehension barriers [3]. Indeed, many users, especially those with low health literacy [4], experience difficulty performing tasks related to health interpretation when using PHRs [5],[6]. Many improvements have been suggested that increase patients’ abilities to understand their personal health information more holistically, such as using graphs to track trends in clinical measurements, e.g. blood pressure or blood glucose levels [7],[8],[9].

To reduce comprehension barriers related to the complexity of medical jargon and interpretation of clinical data, we have synthesized patients’ clinical data into a single quantifiable value using clinically validated risk scores. For example, the Reynolds risk score [10],[11] was previously visualized (https://apps.smarthealthit.org/app/cardiac-risk) by showing patients their blood test results along with a scale of desirable to high risk of cardiovascular disease. This example displays a patient’s cardiac risk in a way that was intended to be more interpretable to those without a medical background. The visualization is able to use a patient’s personal data through a secure EHR technology, SMART on FHIR [12]. We complement and expand upon such existing solutions by designing an application that not only presents a patient with their risk scores but also provides details about clinical measurements and external factors (e.g., pollen count) in a comprehensive patient portal that allows a patient to actively engage with their own data.

Our application introduces visualizations that allow patients to interpret clinical measurements to improve their health understanding. Further, we have created visualizations of patients’ longitudinal data, which gives them the ability to compare their current health risks with their risk in the past and their hypothetical risk in the future. This comparison can provide patients insights into the way their health has changed over time. By creating a resource that allows patients to engage with and explore their health data, we aim to increase patients’ understanding of their health, stimulate the use of preventive health services, encourage patients to initiate informed conversations with their physicians, and improve patient outcomes; previous work has shown that PHRs have helped patients become aware of relevant preventative health services and have more effective conversations with their physicians [13].

## METHODS

### Risk Scores

Clinical measurements from a sample Fast Health Interoperability Resources (FHIR) (https://www.hl7.org/fhir/index.html) data store (https://fhirtest.uhn.ca/about) were synthesized. Twenty-eight risk scores were collected from a review of the literature and five were selected (Table 1) to be used for this proof-of-concept, based on their relative applicability to a variety of patients, their clinical validity, and availability of necessary clinical variables available in the sample EHR. Clinical variables (e.g., age, body mass index, blood pressure, etc.) are extracted from the FHIR database, that serves as the proxy for an actual EHR. While these risk scores may not necessarily be relevant to every patient, they demonstrate the broad applicability of our application. We have included a section in the measurements panel for patients to enter data not conventionally stored accurately in the EHR, such as family history of a certain disease or smoking status, as some risk scores require these data.

**Table 1.** Risk calculators implemented in Halyos.

| Disease Referenced | Measurements Used |
| --- | --- |
| Cardiovascular Disease (Reynolds Risk Score) [20,21] | Gender; Smoker; BP; Cholesterol; HDL; hsCRP; family history |
| Non-alcoholic fatty liver disease [22] | Age; Body Mass Index; Presence of impaired fasting glucose or diabetes; Aspartate aminotransferase/Alanine aminotransferase ratio; Platelet count; Albumin |
| Kidney Failure from Chronic Kidney Disease [23] | Age; Sex; GFR; Albumin creatinine |
| Mortality Risk from Chronic Obstructive Pulmonary Disorder [24] | Blood Urea Nitrogen; Mental status; Heart rate; Age |
| Risk of developing diabetes [25] | Age; Body Mass Index; Waist Circumference; History of antihypertensive drug treatment; high blood glucose; gender |

### Implementation

The design of the user interface was informed by a human-centered design workshop involving patient and clinician input. It was then developed with front-end frameworks, React.js (https://reactjs.org/) and Redux (https://redux.js.org/). There is strong evidence that a clean user interface is valuable, as patient portal usability has been shown to play a role in patient engagement [14]. All data visualization components were built using D3 [15] and custom HTML/JS components which enable the creation of responsive visualizations for patient interaction. The source code of Halyos is available at https://github.com/hms-dbmi/halyos/.

### SMART on FHIR

Substitutable Medical Applications and Reusable Technologies (SMART) on Fast Healthcare Interoperability Resources (FHIR) is a platform that allows 3rd party applications to securely send requests to EHR systems to retrieve patient data [12]. We utilized the SMART on FHIR platform during development to securely enable the integration of personalized patient data.

## RESULTS

### Risk Scores

When a patient first uses Halyos, the most prominent features are the risk scores located at the top (Figure 1, Label 1). Because all measurements required for a risk score may not have been taken at the same time, we calculate present risk scores based on the most recent of each of the measurements required. Upon clicking on a risk score, a section is displayed with more in-depth information (Figure 1, Label 2). Here we provide a risk visualization that displays a patient’s past, present, and hypothetical future risk score as a pictorial unit chart of 100 dots (Figure 1, Label 3). A black dot corresponds to one percentage point of the risk score, e.g., 5 dots indicate 5% risk. Dots of smaller size indicate a decrease compared to the patient’s present risk score. Dots with an outline indicate an increase compared to the present score. This visualization demonstrates how a patient’s risk has changed over time, with particular emphasis on changes from their present value. Pictorial unit charts have been hypothesized to be effective at communicating risks in healthcare settings [16]. By selecting a past date as a point of comparison with their present health, patients can explore how their health has changed over time. To aid with interpretability, only the measurements that contribute to certain risk scores are shown. This filtering allows exploration of the effect of changing the potential future value of their clinical measurements by updating the associated risks as they change their potential future values of relevant clinical measurements.

### Clinical Measurements

The bottom left of the dashboard lists a patient’s clinical measurements (Figure 1, Label 4). It includes the section for patients to enter relevant data that is not typically stored in the EHR. An arrow is included to highlight how clinical measurements have changed since a user-specified past date. Clicking on a measurement reveals a visualization of longitudinal data. This visualization shows the patient’s data of the selected clinical measurement over time and includes shading to represent typical reference ranges, as reported in the EHR (Figure 2, Label 1). Patients can horizontally pan, zoom, and brush the graph to select a time period of interest. The future value is interactively adjustable to help the patient understand the impact of a potential future measurement on various risk scores. Along with this visualization, text is displayed explaining the measurement. For the purpose of this paper, we did not curate audience-appropriate definitions, but this can be done in a clinical setting.

**Figure 2.**
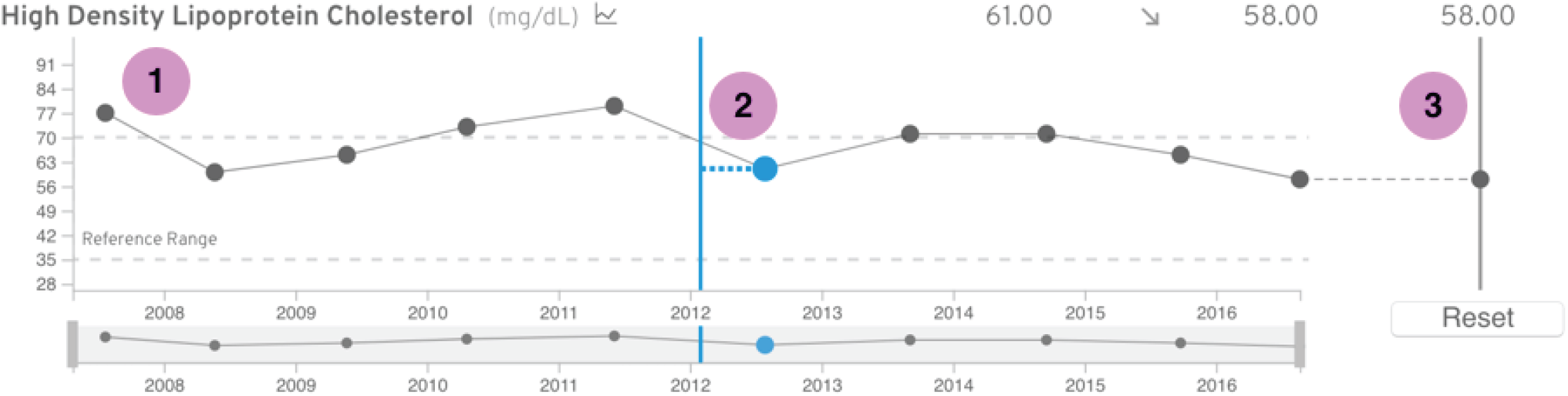
Graph of a patient’s specific measurement values over time (1). The horizontal blue line represents the patient selected past date. The blue dot indicates which value is closest to that past date, and therefore which value is being used to calculate the past risk score (2). The slider on the right allows a patient to manipulate their “future” value, which is used to calculate the future risk score (3).

### Clinical Measurement Visualization

The patient’s ability to select a past date, which is used to find the nearest measurements and calculate “past” risk scores, enables a patient to explore their health data over time. This past date is also displayed on the graph. The measurement used for risk score calculations is displayed as a light blue dot (Figure 2, Label 2). Further temporal exploration is enabled by the previously described ability to manipulate future values. These interactions allow patients to understand how clinical measurements impact multiple risk measures. For example, observing a decrease in Reynold’s risk score as blood pressure declines builds intuition that blood pressure is important for a healthy cardiovascular system.

### Customization

To change or add a new risk score, a clinician would specify a function and indicate (through LOINC codes) which measurements are necessary to calculate it. While Halyos implements five generally applicable risk scores as a proof of concept, the system has been developed to enable the simple addition of more risk scores as long as the risk score-relevant measurements are available in the EHR.

### External Data Integration

We used the US Preventive Services Task Force API to integrate suggested preventive care measures customized to a patient’s gender and age. These suggestions have been shown to increases patient engagement with PHRs, as well as utilization of these preventive services [17]. A number of other non-EHR data sources are integrated to provide patients with a comprehensive view of their health. Location information is used to determine pollen levels, air quality, and flu cases given a geographical region. Pollen levels are taken from the Accuweather API (https://developer.accuweather.com/), air quality from IQAir (https://airvisual.com/api), and flu cases from Flu Near You by HealthMap [18]. This specific information was chosen to be included based on conclusions from the human-centered design workshop [19].

## DISCUSSION

One of the key contributions of Haylos is the way it presents health data in context, which can potentially lead a patient to a greater understanding of their own health. The temporal aspect of the clinical measurement visualization further contextualizes clinical data, as it allows patients to see changes in health over time. Viewing their past and future disease risks, and the contributing measurements, allows patients to gain an intuition for how certain measurements may impact their health. Our application also allows non-clinical measurements included from external APIs or manually entered data to be used in risk scores, making it powerfully customizable to unique patient populations. Finally, we begin to integrate auxiliary information beyond medical measurements, such as environmental data and custom preventive care suggestions. Unsolved challenges of our current approach include restriction to EHR resources as our only source of patient data. There are important data points not captured within an EHR that can determine the trajectory of a person’s health, such as exercise habits, eating habits, and drug use. While we have included a place for user input of relevant non-EHR data, the level of patient engagement remains to be explored. Additionally, since EHR data is not collected at standard intervals, data categorized together as “present” measurements may actually have been measured months apart. While we present the patient with information about how recent a measurement is, risk scores do not factor in these differences; the result of using temporally sparse data may mislead patients.

The design of Halyos addresses concerns raised by clinicians and patients in various studies, such as difficulty understanding medical terminology and lack of appropriate visualization [5] [8]. Future work will involve a user study to comprehensively assess the effectiveness of the design and to provide deeper links between data source, such as customizing preventive care suggestions within the context of the environment (e.g., suggesting a user to exercise indoors if outdoor air quality is poor). Developing an administrative user interface would allow clinicians to customize new risk scores to a particular patient. The eventual goal is to integrate Halyos into a live EHR system to test its efficacy in increasing patient health.

## CONCLUSION

Our work contributes representations of clinical data that aim to make a patient’s health more interpretable through both data synthesis via risk scores and the ability to temporally explore these personalized health data. As tools like SMART on FHIR improve patients’ accessibility to their personal health data, it becomes of utmost importance to develop meaningful ways for patients to explore and interact with their health. To this end, we developed Halyos to aggregate and synthesize patient data, and visualize it in a way that encourages exploration. This exploration was designed to allow patients to gain intuition about their health, and stimulate conversation with physicians, resulting in patients making more informed decisions about their care. This work is particularly timely, as interoperability increasingly allows for personalized medicine solutions to easily adapt to a patient’s data.

## ACKNOWLEDGEMENTS

The authors acknowledge member of the Gehlenborg Lab and Susanne Churchill for mentorship and support. We would also like to acknowledge the participants of our user-centered design workshop. This work was enabled by NIH grants U54HG007963 and R00HG007583.

## REFERENCES

1 Sarkar U, Karter AJ, Liu JY, et al. The Literacy Divide: Health Literacy and the Use of an Internet-Based Patient Portal in an Integrated Health System—Results from the Diabetes Study of Northern California (DISTANCE). Journal of Health Communication. 2010; 15 :183–96. doi:10.1080/10810730.2010.499988

2 Choi NG, Dinitto DM. The digital divide among low-income homebound older adults: Internet use patterns, eHealth literacy, and attitudes toward computer/Internet use. J Med Internet Res 2013; 15 :e93.

3 Goldzweig CL, Orshansky G, Paige NM, et al. Electronic Patient Portals: Evidence on Health Outcomes, Satisfaction, Efficiency, and Attitudes. Annals of Internal Medicine. 2013; 159 :677. doi:10.7326/0003-4819-159-10-201311190-00006

4 Baker DW. The meaning and the measure of health literacy. Journal of General Internal Medicine. 2006; 21 :878–83. doi:10.1111/j.1525-1497.2006.00540.x

5 Tieu L, Schillinger D, Sarkar U, et al. Online patient websites for electronic health record access among vulnerable populations: portals to nowhere? J Am Med Inform Assoc 2017; 24 :e47–54.

6 Tieu L, Sarkar U, Schillinger D, et al. Barriers and Facilitators to Online Portal Use Among Patients and Caregivers in a Safety Net Health Care System: A Qualitative Study. J Med Internet Res 2015; 17 :e275.

7 Sox CM, Gribbons WM, Loring BA, et al. Patient-centered design of an information management module for a personally controlled health record. J Med Internet Res 2010; 12 :e36.

8 Haggstrom DA, Saleem JJ, Russ AL, et al. Lessons learned from usability testing of the VA’s personal health record. Journal of the American Medical Informatics Association. 2011; 18 :i13–7. doi:10.1136/amiajnl-2010-000082

9 Britto MT, Jimison HB, Munafo JK, et al. Usability testing finds problems for novice users of pediatric portals. J Am Med Inform Assoc 2009; 16 :660–9.

10 Ridker PM, Buring JE, Rifai N, et al. Development and validation of improved algorithms for the assessment of global cardiovascular risk in women: the Reynolds Risk Score. JAMA 2007; 297 :611–9.

11 Ridker PM, Paynter NP, Rifai N, et al. C-Reactive Protein and Parental History Improve Global Cardiovascular Risk Prediction. Circulation. 2008; 118 :2243–51. doi:10.1161/circulationaha.108.814251

12 Mandel JC, Kreda DA, Mandl KD, et al. SMART on FHIR: a standards-based, interoperable apps platform for electronic health records. J Am Med Inform Assoc 2016; 23 :899–908.

13 Ant Ozok A, Wu H, Garrido M, et al. Usability and perceived usefulness of Personal Health Records for preventive health care: a case study focusing on patients’ and primary care providers’ perspectives. Appl Ergon 2014; 45 :613–28.

14 Irizarry T, DeVito Dabbs A, Curran CR. Patient Portals and Patient Engagement: A State of the Science Review. J Med Internet Res 2015; 17 :e148.

15 Bostock M, Ogievetsky V, Heer J. D 3 : Data-Driven Documents. IEEE Trans Vis Comput Graph 2011; 17 :2301–9.

16 Edwards A. Explaining risks: turning numerical data into meaningful pictures. BMJ. 2002; 324 :827–30. doi:10.1136/bmj.324.7341.827

17 Krist AH, Woolf SH, Bello GA, et al. Engaging primary care patients to use a patient-centered personal health record. Ann Fam Med 2014; 12 :418–26.

18 Chunara R, Aman S, Smolinski M, et al. Flu Near You: An Online Self-reported Influenza Surveillance System in the USA. Online J Public Health Inform 2013; 5. doi:10.5210/ojphi.v5i1.4456

19 Kerzner E, Goodwin S, Dykes J, et al. A Framework for Creative Visualization-Opportunities Workshops. IEEE Trans Vis Comput Graph Published Online First: 20 August 2018. doi:10.1109/TVCG.2018.2865241

20 Ridker, P. M., Buring, J. E., Rifai, N., & Cook, N. R. (2007). Development and validation of improved algorithms for the assessment of global cardiovascular risk in women: the Reynolds Risk Score. Jama, 297(6), 611–619.

21 Ridker PM, Paynter NP, Rifai N, Gaziano JM, Cook NR. C-reactive protein and parental history improve global cardiovascular risk prediction: the Reynolds Risk Score for men. Circulation. 2008 Nov 25;118(22):2243–51.

22 Angulo P, Hui JM, Marchesini G, Bugianesi E, George J, Farrell GC, Enders F, Saksena S, Burt AD, Bida JP, Lindor K. The NAFLD fibrosis score: a noninvasive system that identifies liver fibrosis in patients with NAFLD. Hepatology. 2007 Apr 1;45(4):846–54.

23 Tangri N, Stevens LA, Griffith J, Tighiouart H, Djurdjev O, Naimark D, Levin A, Levey AS. A predictive model for progression of chronic kidney disease to kidney failure. Jama. 2011 Apr 20;305(15):1553–9.

24 Shorr AF, Sun X, Johannes RS, Yaitanes A, Tabak YP. Validation of a novel risk score for severity of illness in acute exacerbations of COPD. Chest 2011;140(5):1177–1183

25 Lindström J, Tuomilehto J. The diabetes risk score: a practical tool to predict type 2 diabetes risk. Diabetes care. 2003 Mar 1;26(3):725–31.

